# EEG spectral power, but not theta/beta ratio, is a neuromarker for adult ADHD

**DOI:** 10.1101/700005

**Authors:** Hanni Kiiski, Marc Bennett, Laura M. Rueda-Delgado, Francesca Farina, Rachel Knight, Rory Boyle, Darren Roddy, Katie Grogan, Jessica Bramham, Clare Kelly, Robert Whelan

**Affiliations:** Trinity College Institute of Neuroscience, Trinity College Dublin, Dublin, Ireland; UCD School of Psychology, University College Dublin, Dublin, Ireland; Medical Research Council-Cognition and Brain Sciences Unit, University of Cambridge, UK; Global Brain Health Institute, Trinity College Dublin, Dublin, Ireland; Department of Psychiatry, Trinity College Institute of Neuroscience, Trinity College Dublin, Ireland; Department of Physiology, School of Medicine, University College Dublin, Dublin 4, Ireland

**Keywords:** Adults, Attention-Deficit/Hyperactivity Disorder, Classification, Endophenotype, Machine learning, Resting-state EEG

## Abstract

**Background:** Adults with attention-deficit/hyperactivity disorder (ADHD) have been described as having altered resting-state electroencephalographic (EEG) spectral power and theta/beta ratio (TBR). However, a recent review (Pulini et al. 2018) identified methodological errors in neuroimaging, including EEG, ADHD classification studies. Therefore, the specific EEG neuromarkers of adult ADHD remain to be identified, as do the EEG characteristics that mediate between genes and behavior (mediational endophenotypes).

**Methods:** Resting-state eyes-open and eyes-closed EEG were measured from 38 adults with ADHD, 45 first-degree relatives of people with ADHD and 51 unrelated controls. A machine learning classification analysis using penalized logistic regression (Elastic Net) examined if EEG spectral power (1-45 Hz) and TBR could classify participants into ADHD, first-degree relatives and/or control groups. Random-label permutation was used to quantify any bias in the analysis.

**Results:** Eyes-open absolute and relative EEG power distinguished ADHD from control participants (area under receiver operating characteristic = .71-.77). The best predictors of ADHD status were increased power in delta, theta and low-alpha over centro-parietal regions, and in frontal low-beta and parietal mid-beta. TBR did not classify ADHD status. Elevated eyes-open power in delta, theta, low-alpha and low-beta distinguished first-degree relatives from controls (area under receiver operating characteristic = .68-.72), suggesting that these features may be a mediational endophenotype for adult ADHD.

**Conclusions:** Resting-state EEG spectral power may be a neuromarker and mediational endophenotype of adult ADHD. These results did not support TBR as a diagnostic neuromarker for ADHD. It is possible that TBR is a characteristic of childhood ADHD.

## Introduction

Attention-Deficit/Hyperactivity Disorder (ADHD) is a neurodevelopmental disorder whose symptoms emerge in childhood but can continue into adulthood in 40-67% of cases (1–4). ADHD affects 1–7% of adults worldwide and is highly heritable (5–7). Diagnosis is based on subjective behavioural criteria measured through structured clinical interviews and questionnaires (DSM5, Conner’s ADHD scales; 3, 8). However, research increasingly focuses on neural systems, with the goal of discovering complementary objective tools to aid ADHD nosology (9).

One approach to investigating neural systems in ADHD is to record electroencephalography (EEG) while a person is not engaged in any specific task (i.e., ‘resting-state’). Resting-state EEG signals can be decomposed into waves that oscillate at different frequencies. These are typically characterised in terms of delta, theta, alpha, beta and gamma frequency bands (1-4 Hz, 4-8 Hz, 8-13 Hz, 13-30 Hz, 30-45 Hz, respectively). The power in a particular frequency band can be expressed in absolute or relative terms, with relative power indexed as a ratio of one band to the total power in all bands or as a ratio of two bands. EEG resting-state power may provide a neural marker of adult ADHD, although the literature is mixed (10–20). For example, some studies report that adults with ADHD have increased power in fronto-central theta (15) and in posterior (14, 21) or central midline theta (22–24) or higher power in delta (21–24), posterior alpha (14, 22) and beta bands (18, 22, 25). However, others report diminished power in delta (16), theta (15, 16), alpha (15, 16, 25) and in beta range (15, 16, 23, 24). There is also evidence for increased power in theta relative to beta (theta/beta ratio or TBR, 15) in adult ADHD. However, a study with a large sample (N=800) reported *decreased* TBR in adults with ADHD (17). Further, age and developmental course have been shown to affect theta power and TBR more than the presence of ADHD (19, 20).

Biological siblings and parents of individuals with ADHD are 2-8 times more likely to have an ADHD diagnosis, compared to people with no family history of ADHD (5, 6). Heritability of self-rated ADHD symptoms is estimated to be 30-40% in adults based on results from twin and family studies, although first-degree relatives often do not present with symptoms at clinical levels (26). *Mediational endophenotype research* aims to identify neurobiological processes mediating the relationship between genes and behavior (27, 28). Electrophysiological measures are well-suited for mediational endophenotype research because they are highly heritable (29, 30).

Recently, machine learning algorithms have used EEG spectral power, including TBR, to generate classification models that distinguish between childhood ADHD and neuro-typical controls (31). In adults, studies report 68% accuracy for relative theta power (32) and 76% accuracy for power in all frequency bands (18). However, Pulini et al. (31) concluded that methodological factors can lead to inflated accuracy estimates in machine-learning studies using neuroimaging data to classify ADHD. Issues that Pulini (31) identified include circular analysis (including all data when selecting features to be used for classification prior to training), lack of out-of-sample testing and small samples. These factors may result in ‘overfitting’; that is, the fit of the model is disproportionally influenced by the idiosyncrasies in data resulting in a failure to generalize to new datasets. The issue of overfitting is especially relevant to EEG, where the number of input variables typically exceeds the sample size (33). Overfitting of classification models can be diminished by applying methods developed for machine learning classification to relatively large sample sizes (34–36), such as penalized regression. The ‘Elastic Net’ (37) is one such method that can accommodate multi-collinearity (i.e., when independent variables in a regression model are correlated) and select a subset of the most predictive variables (i.e., perform feature selection).

The present study sought to identify the EEG spectral features (i.e., from 1-45 Hz and TBR) that can be considered as neuromarkers and/or mediational endophenotypes for adult ADHD by conducting a classification analysis of adults with ADHD, first-degree relatives and/or unrelated controls. Despite the high heritability of spectral EEG measures (10, 26, 38), no study to date has used a machine-learning classification approach to investigate EEG spectral power as a potential endophenotype for adult ADHD. We expected the first-degree relatives to be best classified by EEG spectral features that are similar to those that distinguish adults with ADHD from controls (27). As most resting-state studies in ADHD have examined EEG when the eyes were closed (e.g. 18, 39), we included both eyes-open and eyes-closed EEG to investigate if either yielded better classification performance. Based on previous studies (14–16, 18, 21–25, 32), and in consideration of Pulini’s (31) findings, we predicted that increased delta and theta power would be the best predictors of ADHD status. Due to the mixed nature of the results (14–16, 18, 21–25, 31, 32) we had no firm hypotheses for alpha, beta and gamma power. Based on Loo et al. (17), we predicted that increase in TBR would not be a good predictor of group membership.

## Methods and Materials

### Participants

Thirty-eight people with an ADHD diagnosis (50% female), 45 first-degree relatives of people with ADHD (18 siblings, 27 parents; 77.8% female) and 51 controls with neither an ADHD diagnosis nor a first-degree relative with an ADHD diagnosis (56.9% female). Participants were recruited through a targeted advertisement campaign to recruit individuals with and without a diagnosis of ADHD as well as family members of individuals with ADHD (Tables 1, S1; see Supplementary Method 1). ADHD group inclusion criteria required the participant to have a formal diagnosis of ADHD *and* a current a T-score over 65 in the ADHD index subscale of Conners’ Adult ADHD Rating Scale (CAARS, 8). The first-degree relative group inclusion criteria required the relative to share both biological parents with, or to be the biological parent of, a person diagnosed with ADHD. For inclusion in the healthy control group, absence of ADHD was required (i.e. no diagnosis of ADHD and a T-score below 65 in the ADHD index subscale of CAARS).

**Table 1.**
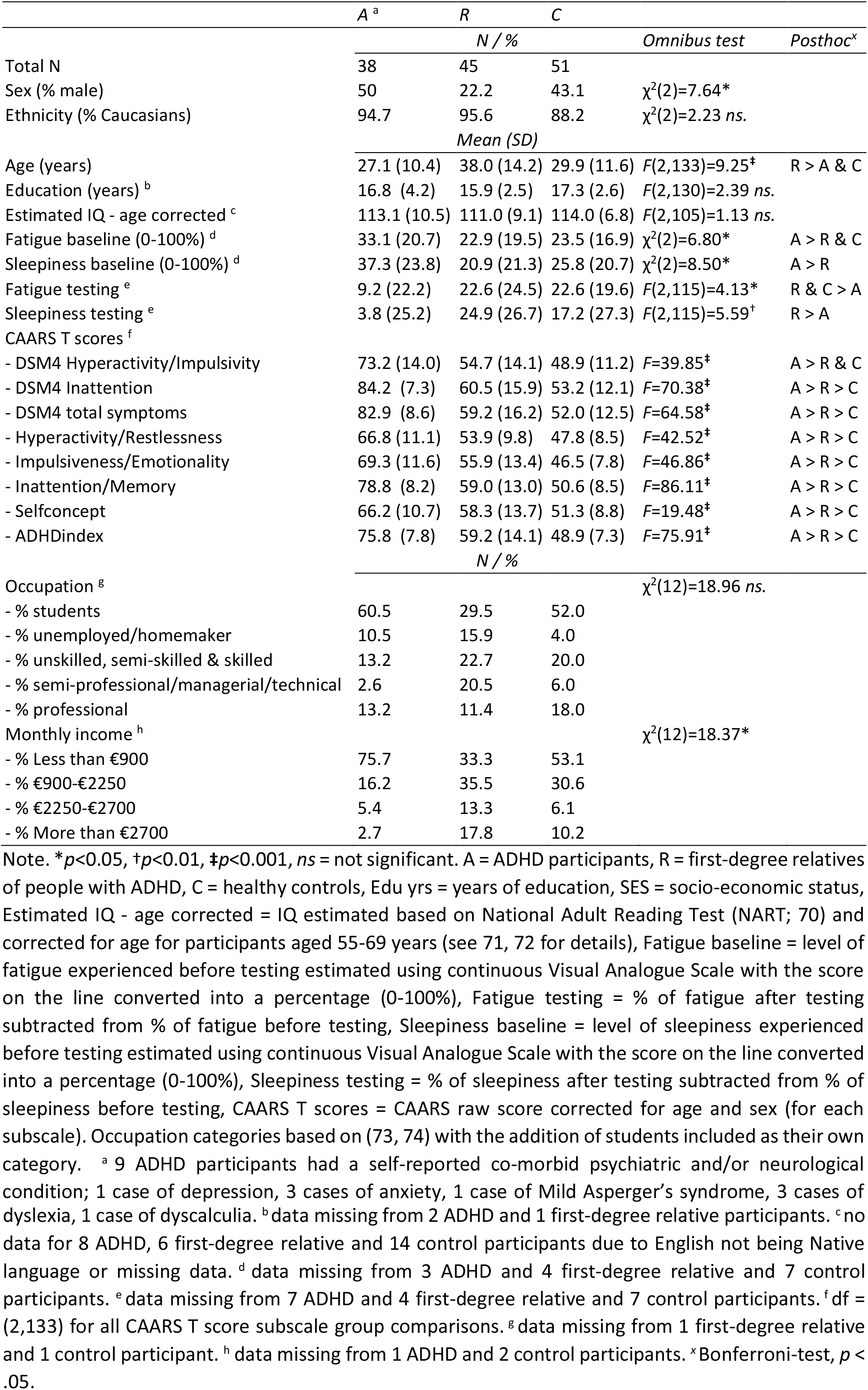
The demographic details and the ADHD symptom profile of the participants.

Participants received €40 euro plus travel expenses. Ethical approval was obtained from the School of Psychology Research Ethics Committee of Trinity College Dublin, the Human Research Ethics Committee of University College Dublin and the Research Ethics Committee of St. Patrick’s Mental Health Services. The research was conducted according to the principles expressed in the Declaration of Helsinki. Written informed consent was obtained from all participants.

### Procedure

Resting-state EEG was collected as part of a larger study on ADHD. Prior to in-laboratory testing, participants completed a battery of online surveys, including demographic details and the CAARS (8). ADHD participants taking methylphenidate (N=14) or lisdexamfetamine (N=3) abstained from medication for at least 36 hrs prior to EEG data collection. On the day of testing, participants were screened for co-morbidities using a shortened version of the Structural Clinical Interview DSM-IV. Full recruitment criteria and screening methods are described in Supplementary Method 1.

Participants were seated in a quiet, darkened room for EEG testing. EEG data were recorded using the ActiveTwoBiosemi™ system from 70 electrodes (64 scalp electrodes, www.biosemi.com), organized according to the 10–5 system (40). Vertical and horizontal electro-oculograms were recorded bilaterally from approximately 2cm below the eye and from the outer canthi, respectively. Two electrodes were placed on the mastoids. Participants were seated 1.05m from a cathode ray tube computer monitor with screen resolution of 1024×768 pixels and a refresh rate of 75 Hz. Data were collected during eyes-open and eyes-closed conditions, each 3.15 mins duration.

### Data analysis

#### EEG pre-processing and analysis

EEG data pre-processing employed the EEGLAB toolbox (41, http://sccn.ucsd.edu/eeglab) in conjunction with the FASTER plug-in (42, http://sourceforge.net/projects/faster). EEG data were bandpass filtered between 1-95 Hz, with the notch frequency set to 48-52 Hz. FASTER automatically identified and removed artefactual (i.e., non-neural) independent components, removed epochs with large artefacts (e.g., muscle twitch) and interpolated channels with poor signal quality (all FASTER pre-processing parameters are included in ‘Supplementary Material_FASTER processing.mat’). Visual inspection of the EEG data was then performed. The EEG was then epoched into 2-s segments.

Spectral analysis of absolute and relative power for 1-45 Hz was conducted using the multitaper spectral estimation with Hanning taper and 0.5 frequency resolution in SPM12 (www.fil.ion.ucl.ac.uk/spm/software/spm12). For absolute power, the raw power values were rescaled to dB. Relative power was calculated as the ratio of the power at that frequency to the total power across all 45 frequencies. The ratio between theta and beta absolute power (i.e. TBR) was calculated by dividing the average absolute power in the theta band (4-8 Hz) with the average absolute power in the low beta band (13-21 Hz; 13, 43–46). This created a TBR for each scalp location.

The EEG frequency data were subsequently parcellated based on both spatial and frequency domains. Data were averaged into 60 spatial bins (i.e., 8-by-8 grid over the scalp, excluding the 4 corners which likely contained non-neural signals), and across each of the 45 frequencies. The spatial parameters were chosen based on previous research (47, 48) as they allow sufficient differentiation among spatial locations without introducing redundant features. Each participant’s input to each prediction model was 2,700 features for EEG absolute/relative power models. Thus, one feature in a prediction model was the power in one spatio-frequency bin that had a specific location on the scalp for one specific frequency (Hz). There were 60 features per participant for TBR models. The same procedure was performed for both eyes-closed and eyes-open tasks.

#### Machine Learning Analysis

Penalized logistic regression with the Elastic Net (37) was used for classification (as in Refs. 48–50, detailed description in Supplementary Method 2). Briefly, the data were initially divided into ten cross-validation (CV) folds. Each of these folds was comprised of a training set (90% of the data) and an out-of-sample test set (10% of the data). The training set was further split into a nested training set (81% of the data) and a nested test set (9% of the data). The analysis consisted of two steps: feature selection followed by model optimization using the Elastic Net.

In the feature selection step, a logistic regression model was fitted on the nested training set using an individual feature as a predictor. The model was evaluated on the nested test set and that feature’s accuracy was measured with the area under the curve of the receiver-operating characteristic (*AROC*). In the model optimization step, selected features were used as predictors in a penalized logistic regression with a range of Elastic Net parameters on the nested training set and subsequently applied to the nested test set. The combination of model parameters and thresholds that resulted in the model with the highest AROC was identified for each nested CV partition, which were then used to create the final model in each main CV fold. This analysis was conducted ten times, using a different CV fold allocations each time so that every participant was contained in the out-of-sample test set at least once.

The entire analysis was repeated 100 times in order to attenuate idiosyncrasies of any given model. Results are mean values across all iterations. The performance of each model was further validated by contrasting model-fit against that of a null model that was generated using random-label permutation (i.e., participants were randomly assigned to a group).

We measured the prediction accuracy of a model with AROC, sensitivity and specificity. A t-test across the 100 iterations, with a significance threshold of *p*<.01, was conducted to compare the prediction accuracy (AROC) of the null model to the model with real data (i.e., actual model). The following classifications were conducted (1) ADHD vs. controls, (2) first-degree relative vs. control, and (3) ADHD vs. first-degree relative. The machine learning analysis was run separately on datasets containing absolute or relative spectral power, or TBR, from either eyes-open or eyes-closed condition (18 models in total). As first-degree relatives were older than the other two groups, we conducted an additional age-matched analyses on the models first-degree relative vs. control and ADHD vs. first-degree relative (see Supplementary Results, Figure S1, Tables S3-6, Videos 11-15).

## Results

The mean number of epochs was 88.0 (*SD*=8.6, range 51-117) and 85.9 (*SD*=8.7, range 51-96) in eyes-closed and eyes-open, respectively. Prediction accuracies (*AROC*) with sensitivity and specificity for the classification models are displayed in Tables 2 and S2-3. A schematic representation of the models’ best predictors (i.e., predictors included in the final model for a specific frequency band ≥90% of the time) are shown in Figures 1-2 and S1. Here, we refer to the predictors with the following structure: *scalp region* (frontal/central/parietal); *frequency band* (delta/theta/alpha/beta/gamma); and *frequency band subdivision* if relevant (low/mid/high); for example, ‘central-beta-low’.

**Table 2.**
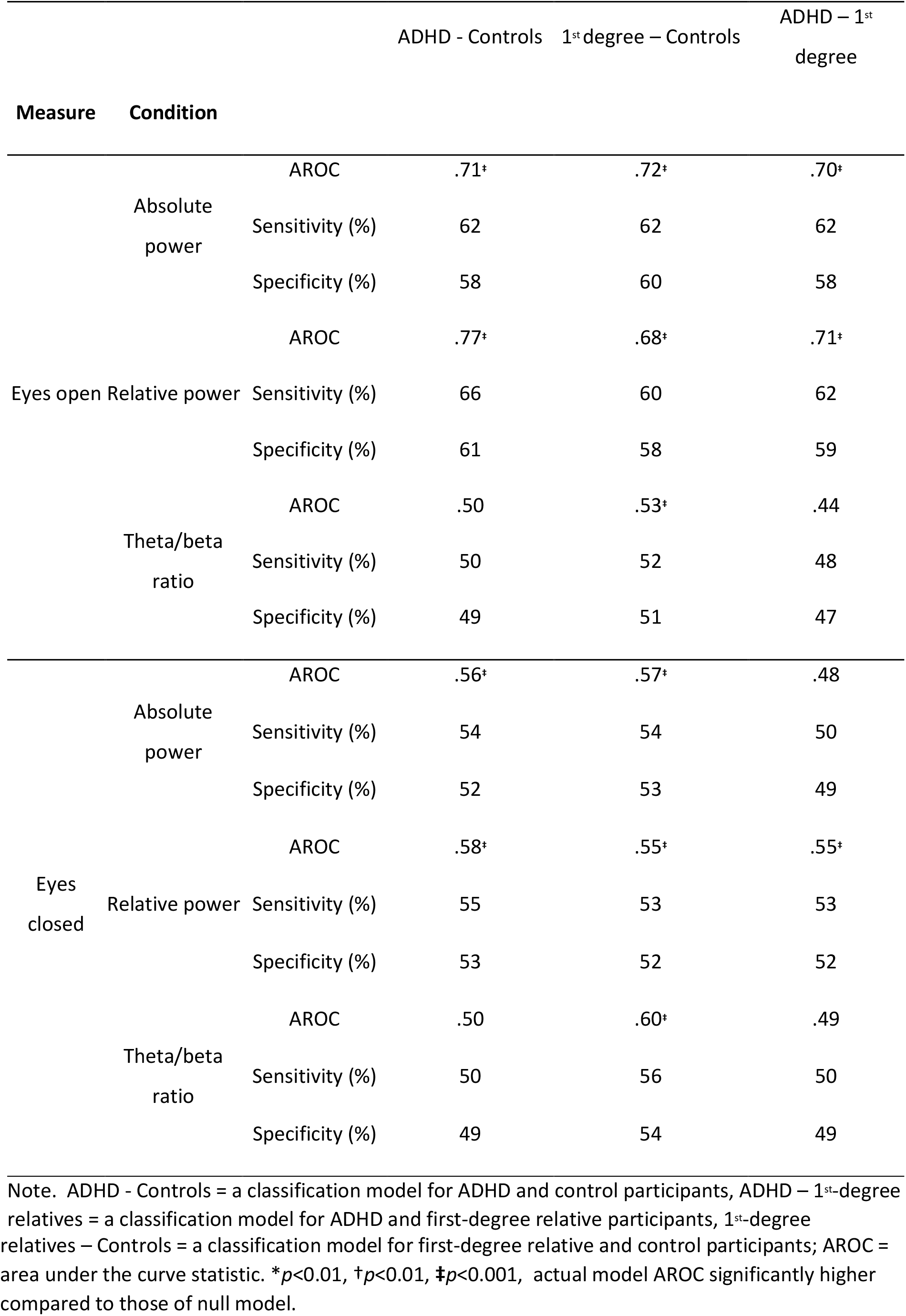
Prediction accuracy, sensitivity and specificity of classification models using eyes-open or eyes-closed resting-state EEG absolute power, relative power and theta/beta ratio as predictors. The mean duration of the final data was 185.2 s for eyes-closed, and 183.7 s for eyes-open. During visual inspection, 3.2% and 2.6% of data was removed in eyes-closed and eyes-open, respectively.

**Figure 1.**
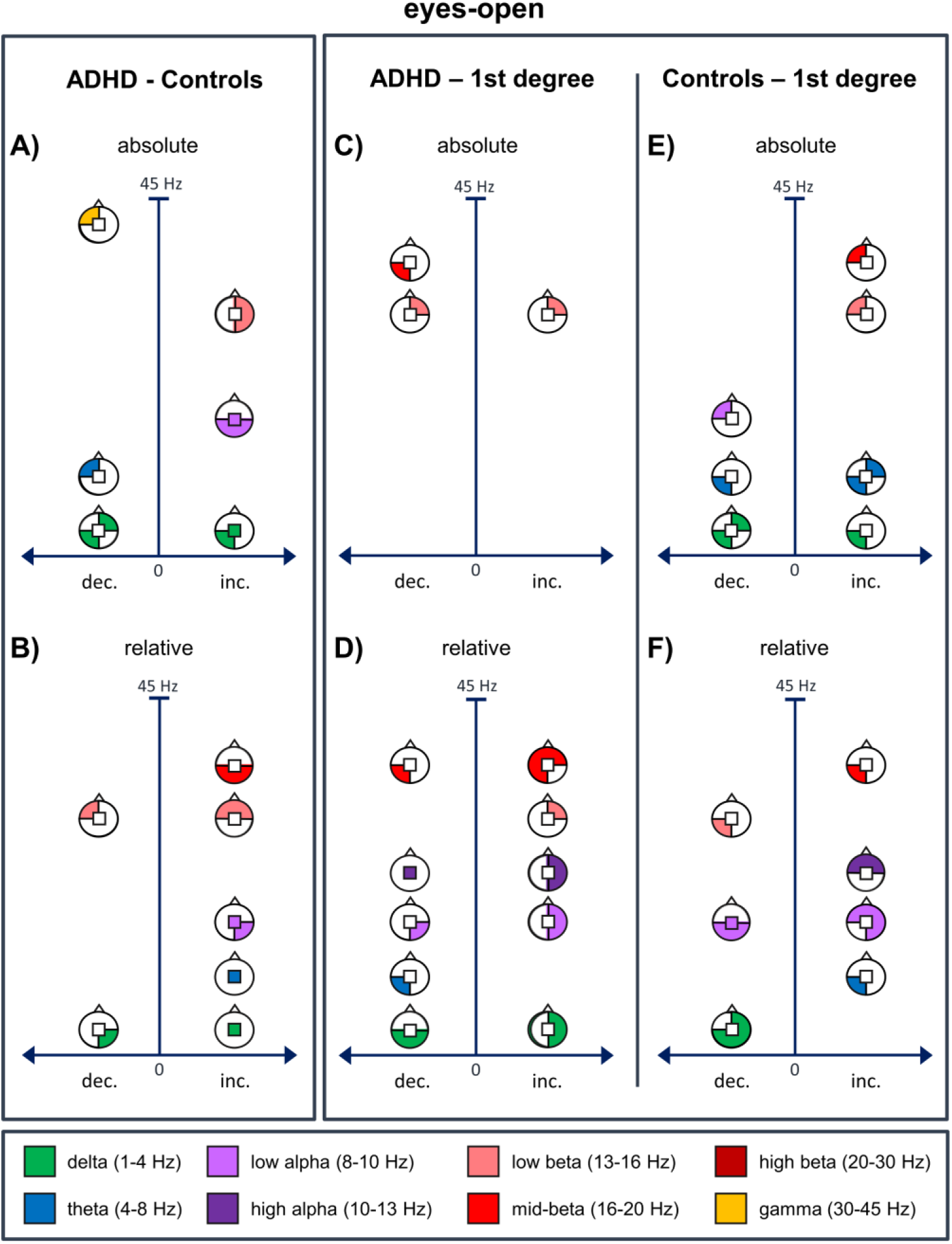
A schematic representation of the best predictors (i.e., predictors included in the final model for a specific frequency band ≥90% of the time) for the eyes-open classification models. 1A) ADHD vs. control model with absolute power (inc./dec. = increase/decrease in power predicts ADHD status), 1B) ADHD vs. control model with relative power (inc./dec. = increase/decrease in power predicts ADHD status), 1C) ADHD vs. first-degree relative model with absolute power (inc./dec. = increase/decrease in power predicts ADHD status), 1D) ADHD vs. first-degree relative model with relative power (inc./dec. = increase/decrease in power predicts ADHD status), 1E) control vs. first-degree relative model with absolute power (inc./dec. = increase/decrease in power predicts first-degree relative status), 1F) control vs. first-degree relative model with relative power (inc./dec. = increase/decrease in power predicts first-degree relative status).

### ADHD vs. control

#### Spectral Power

##### Eyes-open

The AROC for absolute power was .71. The best predictors for ADHD status were increase in central-delta and left-parietal-delta, central-alpha-low and parietal-alpha-low and right-beta low power. The best predictors of control status were increase in right-frontal-delta and left-parietal-delta, left-frontal-theta and gamma power (Figure 1A, Table S4, Video 1). The AROC for relative power was .77. The best predictors for ADHD status were increase in central-delta and central-theta, central-alpha-low and right-parietal-alpha-low, frontal-beta-low and parietal-beta-mid power. The best predictors of control status were increase in right-parietal-delta and left-frontal-beta-low power (Figure 1B, Table S4, Video 2).

##### Eyes-closed

The AROC for absolute power was .56. The best predictors for ADHD status were increase in left-frontal-delta and right-frontal-theta and beta-low power. Elevated right-parietal-alpha-low power was the best predictor for control status (Figure 2A, Table S5, Video 3). The AROC for relative power was .58. The best predictors for ADHD status were increase in left-frontal-delta, left-frontal-theta and right-parietal-theta and parietal-beta-mid power. Increased right-parietal-alpha-low power was the best predictor for control status (Figure 2B, Table S5, Video 4).

**Figure 2.**
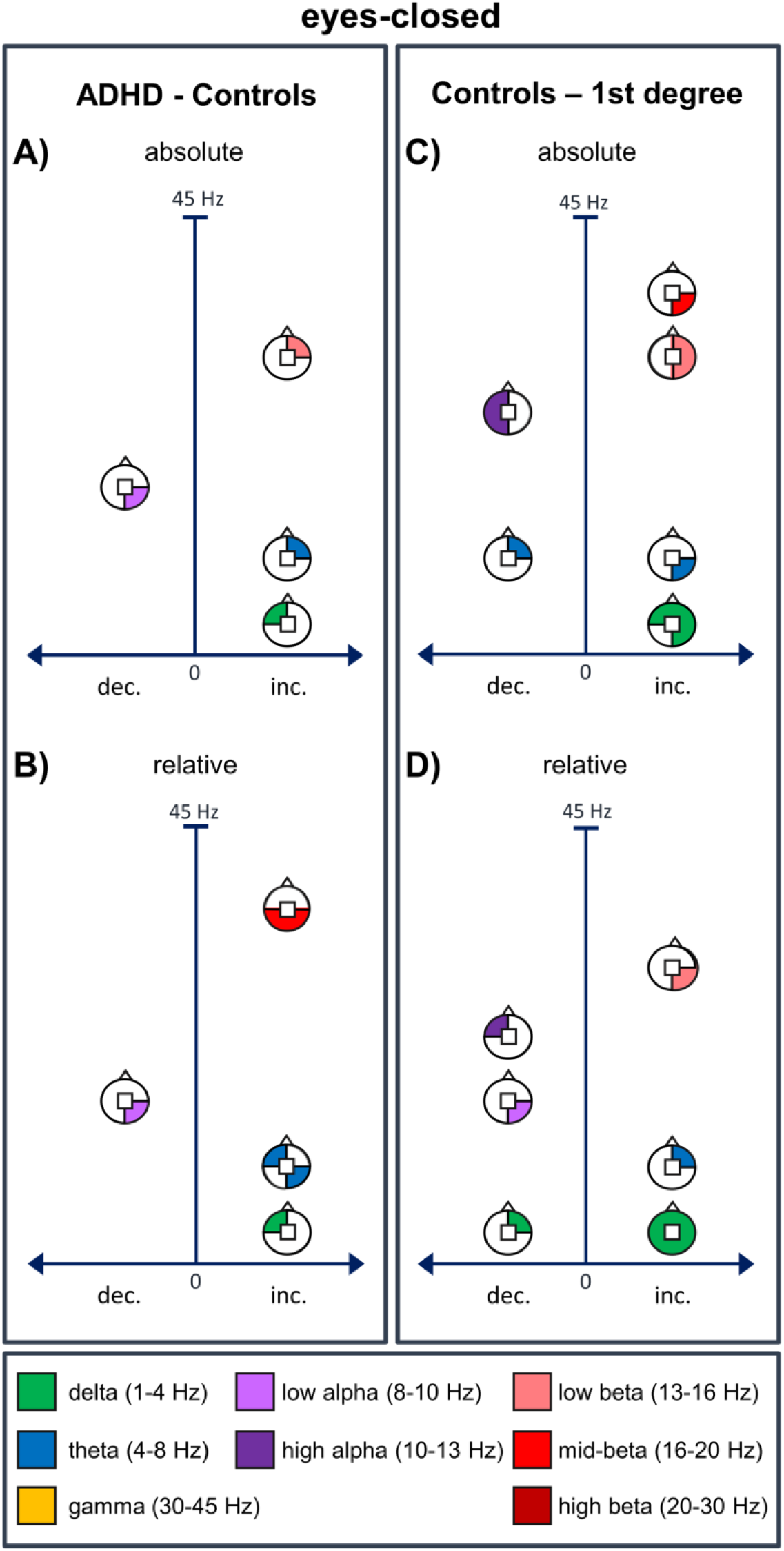
A schematic representation of the best predictors (i.e., predictors included in the final model for a specific frequency band ≥90% of the time) for the eyes-closed classification models. 1A) ADHD vs. control model with absolute power (inc./dec. = increase/decrease in power predicts ADHD status), 1B) ADHD vs. control model with relative power (inc./dec. = increase/decrease in power predicts ADHD status), 1C) control vs. first-degree relative model with absolute power (inc./dec. = increase/decrease in power predicts first-degree relative status), 1D) control vs. first-degree relative model with relative power (inc./dec. = increase/decrease in power predicts first-degree relative status).

#### Theta-beta ratio

Prediction accuracy of TBR was at a chance level in all ADHD classification models (Table 2, S6).

### First-degree relative vs. control

#### Spectral Power

##### Eyes-open

The AROC for absolute power was .72. The best predictors for first-degree relative status were increase in left-parietal-delta, right-frontal-theta and left-parietal-theta and left-frontal-beta-low and left-frontal-beta-mid power. The best predictors for control status were increase in right-frontal-delta and left-parietal-delta, left-parietal-theta and left-frontal-alpha-low power (Figure 1E, Table S4 and Video 5). The AROC for relative power was .68. The best predictors for first-degree relative status were increase in left-parietal-theta, frontal-alpha-low and right-parietal-alpha-low, frontal-alpha-high and left-parietal-beta-mid power. The best predictors for control status were increase in right-frontal-delta and parietal-delta, central-alpha-low and parietal-alpha-low and left-parietal-beta-low power (Figure 1F, Table S4, Video 6).

##### Eyes-closed

The AROC for absolute power was .57. The best predictors for first-degree relative status were increase in right-parietal-delta and frontal-delta, right-parietal-theta, right-beta-low and right-parietal-beta-mid power. The best predictors for control status were elevated power in right-frontal-theta and left-alpha-high (Figure 2C, Table S5, Video 7). The AROC for relative power was .55. The best predictors for first-degree relative status were increased power in frontal-delta and parietal-delta, right-frontal-theta and right-parietal-beta-low. The best predictors for control status were elevated power in right-frontal-delta, right-parietal-alpha-low and left-frontal-alpha-high (Figure 2D, Table S5, Video 8).

#### Theta-beta ratio

The AROC for eyes-open TBR was .53. No TBR feature appeared over 90% of models. The AROC for eyes-closed TBR was .60 and the best predictors for control status were higher TBR located over the right-central region (Table S6).

### ADHD vs. first-degree relative classification

#### Spectral power

##### Eyes-open

The AROC for absolute power was .70. The best predictor for ADHD status was an increase in right-frontal-beta-low power, and elevated right-frontal-beta-low and left-parietal-beta-mid were best predictors for first-degree relative status (Figure 1C, Table S4, Video 9). The AROC for relative power was .71. The best predictors for ADHD status were increase in right-lateralized delta, alpha-low and alpha-high power, and right-frontal-beta-mid, frontal-beta-mid and left-parietal-beta-mid power. The best predictors for first-degree relative status were increase in parietal-delta, left-parietal-theta, right-parietal-alpha-low, central-alpha-high and left-parietal-beta-mid power (Figure 1D, Table S4, Video 10).

##### Eyes-closed

The AROC for relative power was .55. The best predictors for ADHD status were increase in right-frontal-alpha-low, left-frontal-alpha-high and parietal-alpha-high and right-parietal-beta-low power. The best predictors for first-degree relative status were increase in left-parietal-delta, right-frontal-alpha-low and right-parietal-beta-low power. AROCs were at chance level for both absolute and relative power.

#### Theta-beta ratio

The eyes-open and eyes-closed TBR models were not significant (AROCs. 44 and. 49, respectively; Table 2, Table S6).

## Discussion

Characterizing the specific EEG spectral power parameters that best classify ADHD and controls in adults may provide a useful neuromarker to inform diagnosis. We used a sophisticated machine learning approach with Elastic Net regularization (37) including out-of-sample validation to quantify classification model performance for adult ADHD. Our findings revealed moderate prediction accuracy for ADHD vs. control classification models with eyes-open absolute and relative power (AROC=.71 and .77, respectively); however, prediction accuracies for eyes-closed models were only slightly above chance (AROCs=.56-.58). We found that elevated absolute and relative power in several frequency bands could distinguish adults with ADHD from controls.

Markovska-Simoska and Pop-Jordanova (32) reported 68% accuracy for eyes-open relative power in theta band only as a marker of ADHD relative to controls. However, and similar to our findings, adults with ADHD have been shown to have increased power in delta and theta bands (14, 21–24), and also in alpha (14, 22) and beta (18, 22, 25) bands compared to controls. In addition, studies of children with ADHD have found eyes-open delta absolute power (51) and low beta absolute power (13-20Hz) (46) to be the best predictors for ADHD status. These findings together with ours generally suggest that, while resting with eyes open, the absolute and relative power in slow waves over centro-parietal regions (i.e. delta, theta and low-alpha), and faster waves in frontal and parietal regions (i.e. low-beta and mid-beta, respectively) are potential neuromarkers for adult ADHD.

Our secondary focus was to identify mediational endophenotypes from resting-state EEG data; these are subclinical markers of gene expression associated with the disorder and should be overrepresented in first-degree relatives compared to unrelated controls (27, 28). Here, first-degree relatives and controls could be distinguished by their eyes-open absolute and relative power (AROC=.72 and AROC=.68, respectively). We also conducted ADHD vs. first-degree relative classification models that showed high prediction accuracies for eyes-open models of absolute and relative power (AROC=.70 and AROC=.71, respectively), and modest prediction accuracy for age-matched samples (AROC=.55-.60) and relative eyes-closed power.

Previous studies investigating ADHD heritability using EEG spectral power have only provided heritability estimates, and similarities/differences in EEG data between relatives, but specific EEG predictors for first-degree relatives remain under-investigated (28, 52). ADHD and first-degree relative groups shared similar predictors; increased eyes-open absolute power in parietal delta and frontal low-beta bands, and increased eyes-open relative power in centro-parietal theta and low-alpha and parietal mid-beta. Also, the predictors for the age-matched first-degree relative status were similar to ADHD predictors; increased eyes-open absolute power in delta, low-alpha, low-beta, and increased eyes-open relative power in delta, theta, low-alpha and low-beta. These features fulfil the criteria for mediational endophenotypes (27, 28) mediating between genes and behaviour in adult ADHD.

Our findings, together with recent reviews and meta-analyses (19, 31, 53, 54), question the proposed usefulness of TBR as a diagnostic neuromarker for (adult) ADHD (12, 13). It is noteworthy that we did not find TBR to successfully distinguish between ADHD and control adults. Our findings are consistent with non-statistically significant and low accuracy estimates (44-59%) for TBR in empirical and meta-analytic studies from the past 6 years (17, 19, 20, 39, 45, 51), in contrast to moderate to high accuracies (66-88%) in early studies (12, 13, 43, 44) and a small number of recent studies (32, 46, 55). While it is possible that TBR is elevated only in a subgroup of ADHD, neither the characteristics of this subgroup nor the brain state reflected has been determined (19, 39, 54). Differences/inadequacies in methodology and analysis (e.g., overfitting, small samples), and/or heterogeneity of ADHD and control samples may also account for the variability in findings (19, 20, 31). Despite this lack of clarity, TBR recently gained support from FDA as a complementary method for childhood and adolescent ADHD clinical assessment (55, 56).

Overall, our results showed spectral power in eyes-open condition to have greater prediction accuracy compared to eyes-closed power. Most previous research in adults only used the eyes-closed condition (e.g. 18, 39). Prediction accuracies of eyes-closed classification models for ADHD status in this study were significant but very modest (AROC=.56-.58), which is similar to the chance level estimates reported (39), but much lower than 67% (18). Differences between eyes-open and eyes-closed power are likely related to states of vigilance and arousal. For example, opening the eyes during resting-state recording causes global alpha desynchronization (57) and increased skin conductance reflecting heightened arousal (57–59). In fact, people with ADHD have lower arousal levels compared to controls (60, 61), which is one of the most reported theories explaining altered EEG spectral power in ADHD (11, 15, 60–63).

The present study has advantages over previous research. First, we used a machine learning approach with penalized regression and out-of-sample validation to estimate prediction accuracies, thus avoiding many of the pitfalls associated with classification models (31). Second, relatively large adult samples of the three groups underwent high-density resting-state EEG (N=134). We included equal numbers of males and females with ADHD, which is more representative of the adult ADHD population: most previous studies were male only (32, 64) or included <40% female (13, 20, 45, 46, 51, 55). Furthermore, ADHD and control groups were sex-balanced and did not differ in age, education or IQ. Third, we included all 1-45Hz frequencies over the whole scalp area as predictors (similar to 18), unlike many other studies that only included theta and beta power and TBR (13, 20, 39, 45). Lastly, while fMRI resting-state is often used in ADHD classification models (31) EEG has less contraindications than fMRI, is relatively inexpensive and easy to use with a wider range of clinical populations.

Some limitations should be highlighted in the interest of future research. First, the analysis involved only internal cross-validation, not external validation (35). Second, the sample size was at the lower end of the recommended number of participants (over 100-150; 31). However, because we used permutation testing to improve the reliability of significance and variability estimates (31) and all of the models were not significant and/or had high accuracies, the likelihood of inflated accuracy estimates is reduced. Third, although it is unlikely that model accuracy was a consequence of medication (55% ADHD participants medication-naïve, and medicated participants completed at least a 36-hour washout period), fatigue/drowsiness caused by medication withdrawal may have influenced the EEG. Fourth, we did not investigate classification accuracy of EEG spectral power for different ADHD presentations/subgroups as per previous studies (43, 44, 65) due to limited numbers. Furthermore, we did not have sufficient numbers of first-degree relatives to investigate siblings and parents separately, which would have been desirable with respect to age-matching. We could also only recruit 3 ADHD-first-degree relative dyads, thus not being able to calculate heritability estimates for EEG spectral power.

### Conclusion

Stratification of individuals based on their EEG characteristics continues to move beyond group-level comparisons and towards sophisticated methodology to identify prognostic *neuromarkers* (31, 66). The present study is consistent with the ‘big data in psychiatry’ approach that has potential to greatly benefit clinical research and practice (66, 67, 68, 69). Our findings support the development and integration of objective and reliable neurophysiological indicators (e.g. eyes-open absolute and relative power, but not TBR or eyes-closed power), into complementary assessment and prognostic tools for adult ADHD.

## Acknowledgements

We would like to express our gratitude to Mr. Andrea Aleni, Ms. Aoife Sweeney, Ms. Laura Rai and Mr. Ernest Mihelj for their assistance in participant recruitment, and in data collection and management. We would also like to thank HADD Ireland for their valuable support with participant recruitment. The address of the corresponding author Prof. Robert Whelan is Lloyd Building, Trinity College Dublin, Dublin 2, Ireland.

This work was supported by Irish Research Council’s Government of Ireland Postdoctoral Fellowship grant to H. Kiiski (grant number GOIPD/2015/777) and M. Bennett’s contribution was supported by the Irish Research Council’s Government of Ireland Postdoctoral Fellowship grant (grant number GOIPD/2016/617) and the Wellcome Trust (107496/Z/15/Z). This work was also supported by Irish Research Council’s Postdoctoral Enterprise Partnership Scholarship (grant number 207401) to F. Farina, Irish Research Council’s Postgraduate Enterprise Partnership Scholarship (grant number EPSPG/2017/277) to R. Boyle, Irish Research Council’s Government of Ireland Postgraduate Scholarship to K. Grogan (GOIPG/2014/294), the Irish Health Research Board grant to the REDEEM group at Trinity College Institute of Neuroscience and Department of Psychiatry of which D. Roddy is a member (grant number 201651.12553), Pathfinder seed grant from Trinity College Dublin to C. Kelly, and Brain & Behavior Research Foundation’s NARSAD Young Investigator grant (grant number 23599) and a Science Foundation Ireland grant (grant number 16/ERCD/3797) to R. Whelan. The other authors did not have funding to disclose. The study sponsors had no involvement in the collection, analysis and interpretation of data and in the writing of the manuscript.

