## Supplementary material for "EEG spectral power, but not theta/beta ratio, is a neuromarker for adult ADHD": ADHD resting-state EEG neuromarkers Supplementary Material_kiiski et al.docx

**Supplementary Results.** Results of the age-matched eyes-open and eyes-closed classification models.

*Age-matched first-degree relative vs. control.* The AROC for eyes-open absolute and relative power were .63 and .64, respectively. The AROC for eyes-closed TBR was .65. Results are reported in Figures S1C-D, Tables S3-4 and S6, Videos 11-12.

*Age-matched ADHD vs. first-degree relatives.* The AROC for eyes-open absolute and relative power were .60 and .55, respectively. The AROC for eyes-closed relative power and TBR were .57 and .57, respectively. Results are reported in Figures S1A-B and S1E, Tables S3-6, Videos 13-15.

**Figure S1.** A schematic representation of the best predictors (i.e., predictors included in the final model for a specific frequency band ≥90% of the time) for the age-matched eyes-open and eyes-closed classification models. 1A) ADHD vs. first-degree relative model with eyes-open absolute power power (inc./dec. = increase/decrease in power predicts ADHD status), 1B) ADHD vs. first-degree relative model with eyes-open relative power power (inc./dec. = increase/decrease in power predicts ADHD status), 1C) first-degree relative vs. control model with eyes-open absolute power (inc./dec. = increase/decrease in power predicts first-degree relative status), 1D) first-degree relative vs. control model with eyes-open relative power (inc./dec. = increase/decrease in power predicts first-degree relative status), 1E) ADHD vs. first-degree relative model with eyes-closed relative power power (inc./dec. = increase/decrease in power predicts ADHD status).

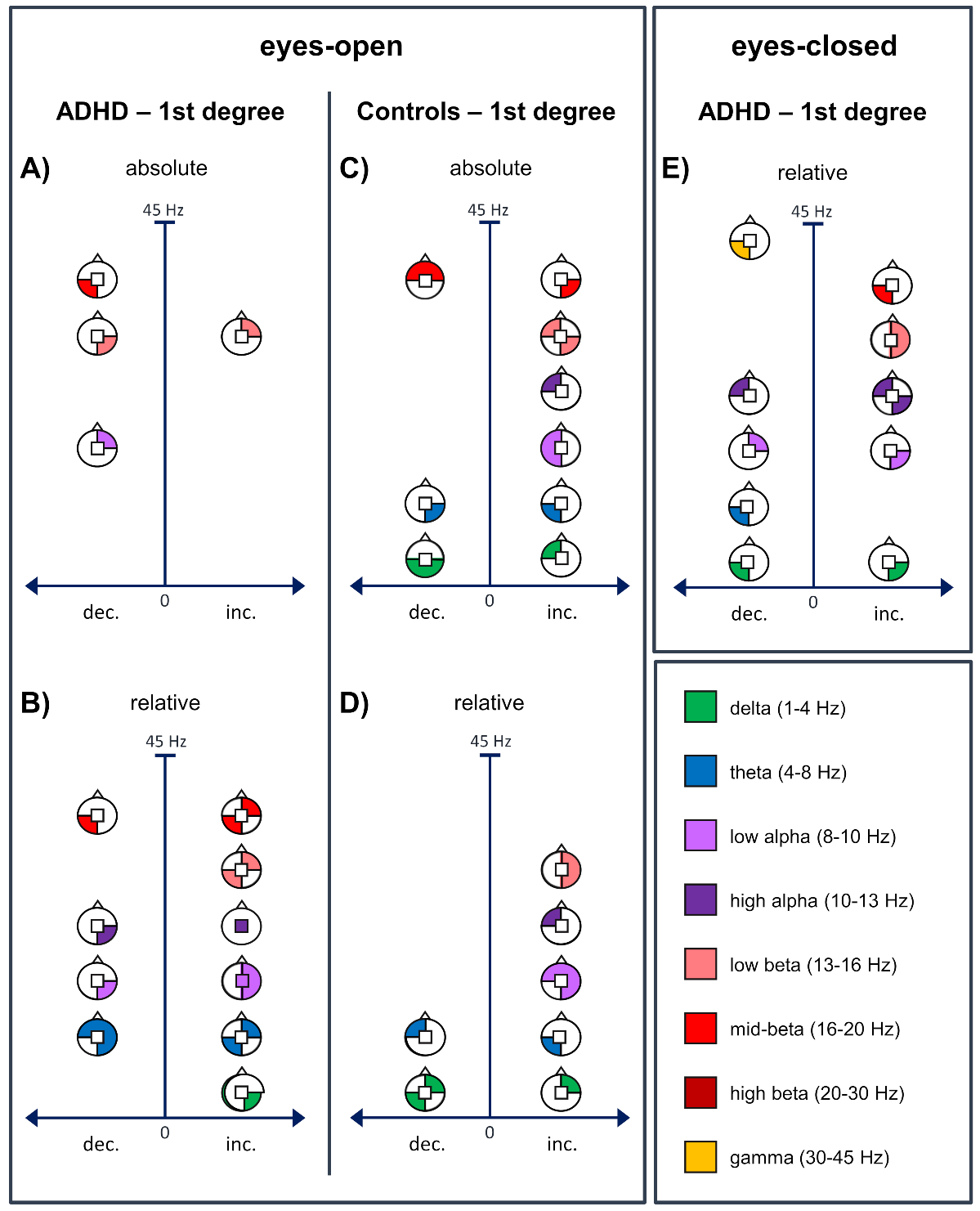

**Table S1.** The full demographic details and the ADHD symptom profile of the participants.

|  | *A* ^a^ | *R* | *C* |  |  |
| --- | --- | --- | --- | --- | --- |
|  | *N / %* | | | *Omnibus test* | *Posthoc^x^* |
| Total N | 38 | 45 | 51 |  |  |
| Sex (% male) | 50 | 22.2 | 43.1 | χ^2^(2)=7.64* |  |
| Ethnicity (% Caucasians) | 94.7 | 95.6 | 88.2 | χ^2^(2)=2.23 *ns.* |  |
|  | *Mean (SD)* | | |  |  |
| Age (years) | 27.1 (10.4) | 38.0 (14.2) | 29.9 (11.6) | *F*(2,133)=9.25**^‡^** | R > A & C |
| Education (years) ^b^ | 16.8 (4.2) | 15.9 (2.5) | 17.3 (2.6) | *F*(2,130)=2.39 *ns.* |  |
| Estimated IQ - age corrected ^c^ | 113.1 (10.5) | 111.0 (9.1) | 114.0 (6.8) | *F*(2,105)=1.13 *ns.* |  |
| Fatigue baseline (0-100%) ^d^ | 33.1 (20.7) | 22.9 (19.5) | 23.5 (16.9) | χ^2^(2)=6.80* | A > R & C |
| Sleepiness baseline (0-100%) ^d^ | 37.3 (23.8) | 20.9 (21.3) | 25.8 (20.7) | χ^2^(2)=8.50* | A > R |
| Fatigue testing ^e^ | 9.2 (22.2) | 22.6 (24.5) | 22.6 (19.6) | *F*(2,115)=4.13* | R & C > A |
| Sleepiness testing ^e^ | 3.8 (25.2) | 24.9 (26.7) | 17.2 (27.3) | *F*(2,115)=5.59^†^ | R > A |
| CAARS T scores ^f^ |  |  |  |  |  |
| - DSM4 Hyperactivity/Impulsivity | 73.2 (14.0) | 54.7 (14.1) | 48.9 (11.2) | *F*=39.85**^‡^** | A > R & C |
| - DSM4 Inattention | 84.2 (7.3) | 60.5 (15.9) | 53.2 (12.1) | *F*=70.38**^‡^** | A > R > C |
| - DSM4 total symptoms | 82.9 (8.6) | 59.2 (16.2) | 52.0 (12.5) | *F*=64.58**^‡^** | A > R > C |
| - Hyperactivity/Restlessness | 66.8 (11.1) | 53.9 (9.8) | 47.8 (8.5) | *F*=42.52**^‡^** | A > R > C |
| - Impulsiveness/Emotionality | 69.3 (11.6) | 55.9 (13.4) | 46.5 (7.8) | *F*=46.86**^‡^** | A > R > C |
| - Inattention/Memory | 78.8 (8.2) | 59.0 (13.0) | 50.6 (8.5) | *F*=86.11**^‡^** | A > R > C |
| - Self concept | 66.2 (10.7) | 58.3 (13.7) | 51.3 (8.8) | *F*=19.48**^‡^** | A > R > C |
| - ADHD index | 75.8 (7.8) | 59.2 (14.1) | 48.9 (7.3) | *F*=75.91**^‡^** | A > R > C |
|  | *N / %* | | |  |  |
| Occupation ^g^ |  |  |  | χ^2^(12)=18.96 *ns.* |  |
| - % students | 60.5 | 29.5 | 52.0 |  |  |
| - % unemployed/homemaker | 10.5 | 15.9 | 4.0 |  |  |
| - % unskilled | 2.6 | 2.3 | 2.0 |  |  |
| - % semi-skilled | 5.3 | 6.8 | 10.0 |  |  |
| - % skilled, clerical/sales | 5.3 | 13.6 | 8.0 |  |  |
| - % semi-professional/managerial/technical | 2.6 | 20.5 | 6.0 |  |  |
| - % professional | 13.2 | 11.4 | 18.0 |  |  |
| Monthly income ^h^ |  |  |  | χ^2^(12)=18.37* |  |
| - % Less than €900 | 75.7 | 33.3 | 53.1 |  |  |
| - % €900-€1350 | 2.7 | 13.3 | 16.3 |  |  |
| - % €1350-€1800 | 5.4 | 11.1 | 8.2 |  |  |
| - % €1800-€2250 | 8.1 | 11.1 | 6.1 |  |  |
| - % €2250-€2700 | 5.4 | 13.3 | 6.1 |  |  |
| - % More than €2700 | 2.7 | 17.8 | 10.2 |  |  |

Note. **p*<0.05, ^†^*p*<0.01, **^‡^***p*<0.001, *ns* = not significant. A = ADHD participants, R = first-degree relatives of people with ADHD, C = healthy controls, Edu yrs = years of education, SES = socio-economic status, Estimated IQ - age corrected = IQ estimated based on National Adult Reading Test (NART; 70) and corrected for age for participants aged 55-69 years (see 71, 72), Fatigue baseline = level of fatigue experienced before testing estimated using continuous Visual Analogue Scale with the score on the line converted into a percentage (0-100%), Fatigue testing = % of fatigue after testing subtracted from % of fatigue before testing, Sleepiness baseline = level of sleepiness experienced before testing estimated using continuous Visual Analogue Scale with the score on the line converted into a percentage (0-100%), Sleepiness testing = % of sleepiness after testing subtracted from % of sleepiness before testing, CAARS T scores = CAARS raw score corrected for age and sex (for each subscale). Occupation categories based on (73, 74) with the addition of students included as their own category. ^a^ 9 ADHD participants had a self-reported co-morbid psychiatric and/or neurological condition; 1 case of depression, 3 cases of anxiety, 1 case of Mild Asperger’s syndrome, 3 cases of dyslexia, 1 case of dyscalculia. ^b^ data missing from 2 ADHD and 1 first-degree relative participants. ^c^ no data for 8 ADHD, 6 first-degree relative and 14 control participants due to English not being Native language or missing data. ^d^ data missing from 3 ADHD and 4 first-degree relative and 7 control participants. ^e^ data missing from 7 ADHD and 4 first-degree relative and 7 control participants. ^f^ df = (2,133) for all CAARS T score subscale group comparisons. ^g^ data missing from 1 first-degree relative and 1 control participant. ^h^ data missing from 1 ADHD and 2 control participants. *^x^* Bonferroni-test, *p* < .05.

**Table S2.** Prediction accuracy, sensitivity and specificity of classification models (actual and null) using eyes-open or eyes-closed resting-state EEG absolute power, relative power and theta/beta ratio (calculated from absolute power) as predictors. The mean duration of the final data was 185.2 s for eyes-closed, and 183.7 s for eyes-open. During visual inspection, 3.2% and 2.6% of data was removed in eyes-closed and eyes-open, respectively.

|  |  |  |  | ADHD - Controls | Controls – first-degree relatives | ADHD – first-degree relatives |
| --- | --- | --- | --- | --- | --- | --- |
| **Measure** | **Condition** | **Model** |  |  |  |  |
| Eyes open | Absolute power | Actual | AROC | .71**^‡^** | .72**^‡^** | .70**^‡^** |
|  |  |  | Sensitivity (%) | 62.40 | 61.98 | 61.62 |
|  |  |  | Specificity (%) | 58.18 | 59.78 | 58.45 |
|  |  | Null | AROC | .51 | .50 | .50 |
|  |  |  | Sensitivity (%) | 50.93 | 50.64 | 50.75 |
|  |  |  | Specificity (%) | 49.69 | 49.58 | 49.50 |
|  | Relative power | Actual | AROC | .77**^‡^** | .68**^‡^** | .71**^‡^** |
|  |  |  | Sensitivity (%) | 66.01 | 60.36 | 61.94 |
|  |  |  | Specificity (%) | 60.84 | 58.32 | 58.70 |
|  |  | Null | AROC | .48 | .50 | .50 |
|  |  |  | Sensitivity (%) | 49.67 | 50.61 | 50.60 |
|  |  |  | Specificity (%) | 48.76 | 49.55 | 49.38 |
|  | Theta/beta ratio | Actual | AROC | .50 | .53**^‡^** | .44 |
|  |  |  | Sensitivity (%) | 50.48 | 52.12 | 47.57 |
|  |  |  | Specificity (%) | 49.36 | 50.91 | 46.89 |
|  |  | Null | AROC | .49 | .49 | .50 |
|  |  |  | Sensitivity (%) | 49.73 | 49.68 | 50.78 |
|  |  |  | Specificity (%) | 48.80 | 48.71 | 49.53 |
| Eyes closed | Absolute power | Actual | AROC | .56**^‡^** | .57**^‡^** | .48 |
|  |  |  | Sensitivity (%) | 54.03 | 54.26 | 49.68 |
|  |  |  | Specificity (%) | 52.02 | 52.69 | 48.59 |
|  |  | Null | AROC | .48 | .50 | .51 |
|  |  |  | Sensitivity (%) | 49.53 | 50.36 | 51.18 |
|  |  |  | Specificity (%) | 48.67 | 49.33 | 49.89 |
|  | Relative power | Actual | AROC | .58**^‡^** | .55**^‡^** | .55**^‡^** |
|  |  |  | Sensitivity (%) | 55.05 | 53.15 | 53.14 |
|  |  |  | Specificity (%) | 52.78 | 51.74 | 51.58 |
|  |  | Null | AROC | .50 | .49 | .49 |
|  |  |  | Sensitivity (%) | 50.84 | 50.13 | 50.02 |
|  |  |  | Specificity (%) | 49.64 | 49.13 | 48.88 |
|  | Theta/beta ratio | Actual | AROC | .50 | .60**^‡^** | .49 |
|  |  |  | Sensitivity (%) | 50.44 | 55.82 | 50.14 |
|  |  |  | Specificity (%) | 49.35 | 54.04 | 48.99 |
|  |  | Null | AROC | .49 | .50 | .49 |
|  |  |  | Sensitivity (%) | 49.85 | 50.71 | 49.91 |
|  |  |  | Specificity (%) | 48.90 | 49.63 | 48.78 |

Note. ADHD - Controls = a classification model for ADHD and control participants; Controls – first-degree relatives = a classification model for control and first-degree relative participants; ADHD – first-degree relatives = a classification model for ADHD and first-degree relative participants; AROC = area under the curve statistic. **p*<0.01, †*p*<0.01, **‡***p*<0.001, actual model AROC significantly higher compared to those of null model.

**Table S3.** Prediction accuracy, sensitivity and specificity of age-matched classification models using eyes-open or eyes-closed resting-state EEG absolute power, relative power and theta/beta ratio as predictors.

|  |  |  |  | Controls –  first-degree relatives | ADHD –  first-degree relatives |
| --- | --- | --- | --- | --- | --- |
| **Measure** | **Condition** | **Model** |  |  |  |
| Eyes open | Absolute power | Actual | AROC | .63**^‡^** | .60**^‡^** |
|  |  |  | Sensitivity (%) | 58.03 | 55.18 |
|  |  |  | Specificity (%) | 54.81 | 54.71 |
|  |  | Null | AROC | .52 | .50 |
|  |  |  | Sensitivity (%) | 51.90 | 50.57 |
|  |  |  | Specificity (%) | 50.14 | 48.98 |
|  | Relative power | Actual | AROC | .64**^‡^** | .55**^‡^** |
|  |  |  | Sensitivity (%) | 58.50 | 53.16 |
|  |  |  | Specificity (%) | 55.17 | 52.20 |
|  |  | Null | AROC | .48 | .49 |
|  |  |  | Sensitivity (%) | 49.49 | 50.46 |
|  |  |  | Specificity (%) | 48.30 | 48.85 |
|  | Theta/beta ratio | Actual | AROC | .49 | .47 |
|  |  |  | Sensitivity (%) | 49.96 | 49.46 |
|  |  |  | Specificity (%) | 48.65 | 47.60 |
|  |  | Null | AROC | .49 | .51 |
|  |  |  | Sensitivity (%) | 50.32 | 51.00 |
|  |  |  | Specificity (%) | 48.93 | 49.52 |
| Eyes closed | Absolute power | Actual | AROC | .44 | .42 |
|  |  |  | Sensitivity (%) | 47.54 | 47.48 |
|  |  |  | Specificity (%) | 46.95 | 44.89 |
|  |  | Null | AROC | .50 | .49 |
|  |  |  | Sensitivity (%) | 50.51 | 50.50 |
|  |  |  | Specificity (%) | 49.09 | 48.88 |
|  | Relative power | Actual | AROC | .47 | .57**^‡^** |
|  |  |  | Sensitivity (%) | 48.92 | 53.74 |
|  |  |  | Specificity (%) | 47.95 | 53.16 |
|  |  | Null | AROC | .50 | .50 |
|  |  |  | Sensitivity (%) | 50.52 | 50.65 |
|  |  |  | Specificity (%) | 49.09 | 49.08 |
|  | Theta/beta ratio | Actual | AROC | .65**^‡^** | .57**^‡^** |
|  |  |  | Sensitivity (%) | 59.71 | 53.69 |
|  |  |  | Specificity (%) | 55.69 | 53.10 |
|  |  | Null | AROC | .50 | .49 |
|  |  |  | Sensitivity (%) | 50.87 | 50.55 |
|  |  |  | Specificity (%) | 49.34 | 48.94 |

Note. Controls – first-degree relatives = a classification model for control and first-degree relative participants; ADHD – first-degree relatives = a classification model for ADHD and first-degree relative participants; AROC = area under the curve statistic. **p*<0.01, †*p*<0.01, **‡***p*<0.001, actual model AROC significantly higher compared to those of null model.

**Table S4.** Beta choice frequencies (all, >90%) and beta values for all the eyes-open classification models. In beta choice frequency plots the scale 0-10 runs from 0=blue to 10=yellow. In beta choice frequency plots >90% yellow denotes frequency-area features over 90%, blue below 90%. In beta value plots green denotes an increase in power and blue a decrease in power in that scalp area to be a predictor for ADHD status in ADHD vs. Controls models. Similarly, green denotes an increase in power and blue a decrease in power in that scalp area to be a predictor for ADHD status in ADHD vs. Relatives model. Finally, green denotes an increase in power and blue a decrease in power in that scalp area to be a predictor for first-degree relative status in Relatives vs. Controls models. These apply also to the age-matched models.

| **Model** |  |  | **Delta** | **Theta** | **low-Alpha** | **high-Alpha** | **low-Beta** | **mid-Beta** | **high-Beta** | **Gamma** |
| --- | --- | --- | --- | --- | --- | --- | --- | --- | --- | --- |
| ADHD vs. Controls | Abs | beta choice freq.  (0-10) | 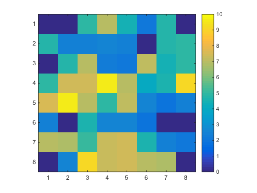 | 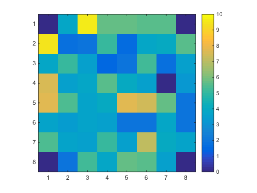 | 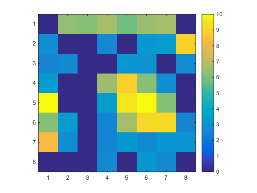 | 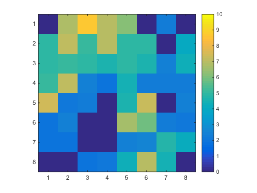 | 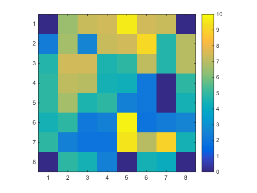 | 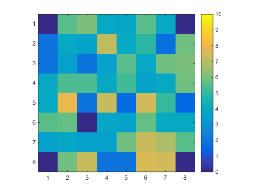 | 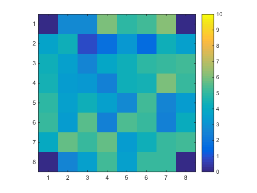 | 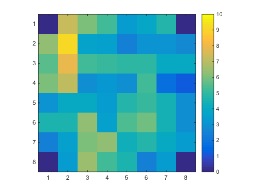 |
|  |  | beta choice freq. >90% | 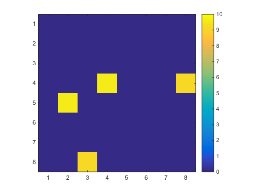 | 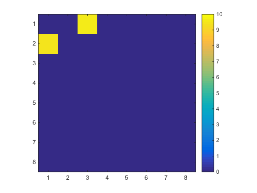 | 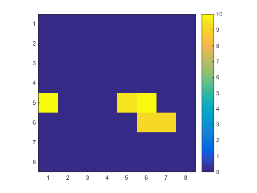 | 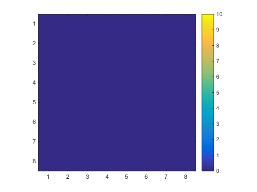 | 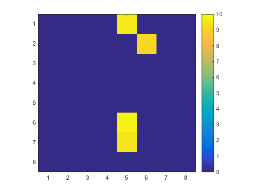 | 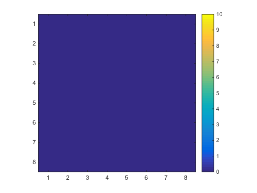 | 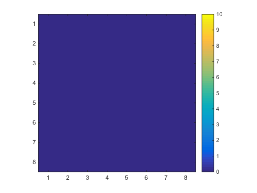 | 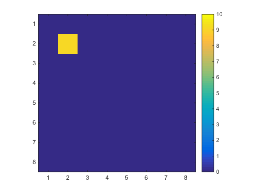 |
|  |  | beta value | 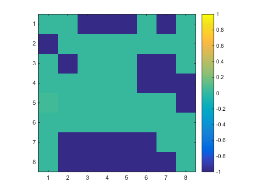 | 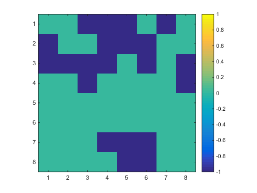 | 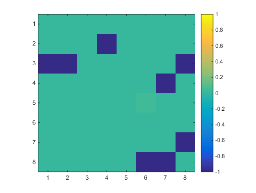 | 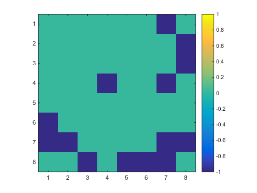 | 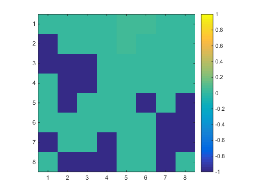 | 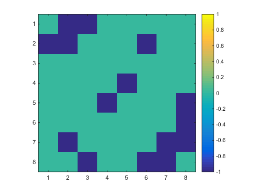 | 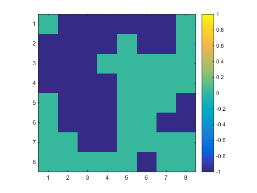 | 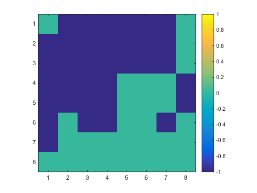 |
|  | Rel | beta choice freq.  (0-10) | 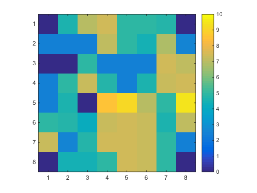 | 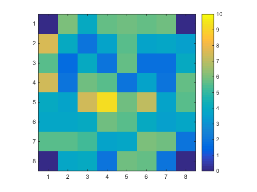 | 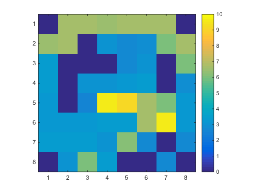 | 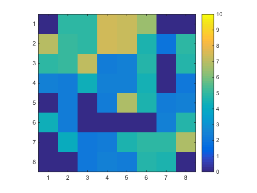 | 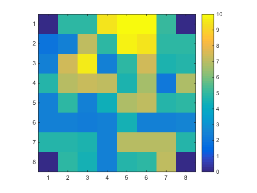 | 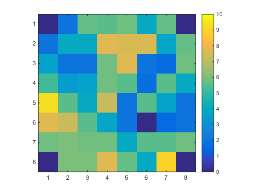 | 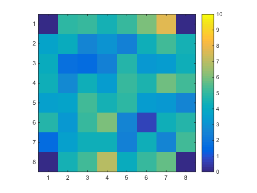 | 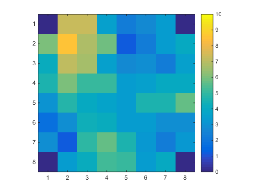 |
|  |  | beta choice freq. >90% | 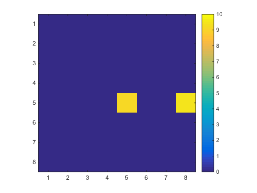 | 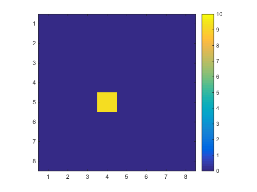 | 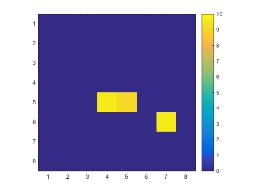 | 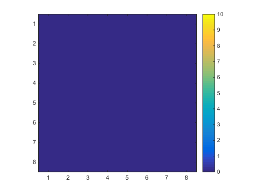 | 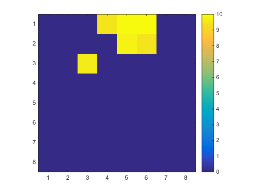 | 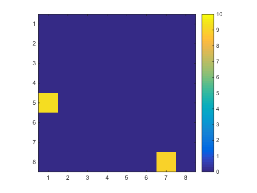 | 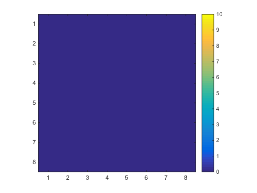 | 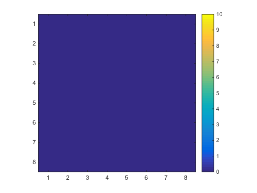 |
|  |  | beta value | 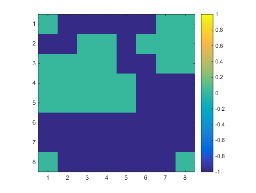 | 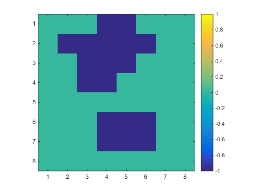 | 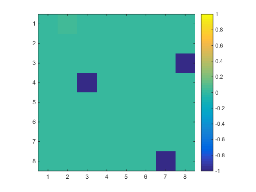 | 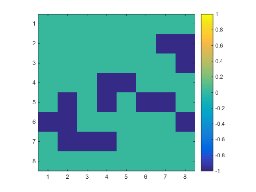 | 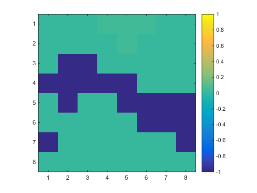 | 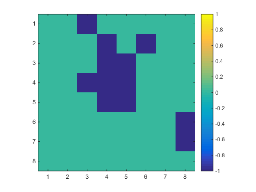 | 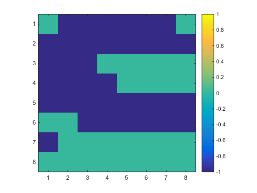 | 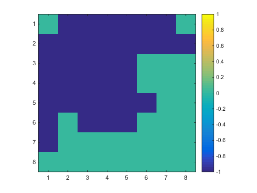 |
| ADHD vs. Relatives | Abs | beta choice freq.  (0-10) | 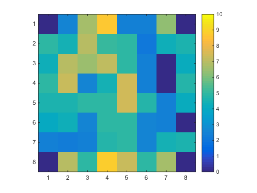 | 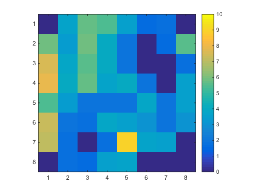 | 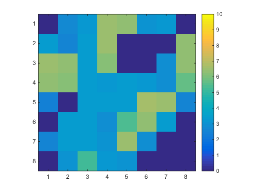 | 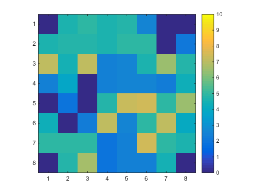 | 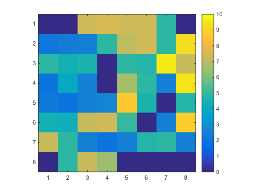 | 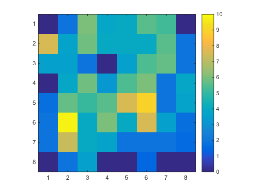 | 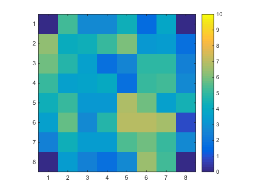 | 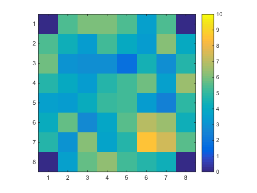 |
|  |  | beta choice freq. >90% | 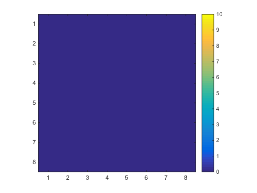 | 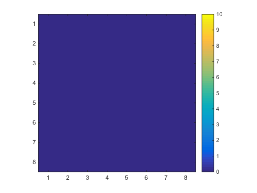 | 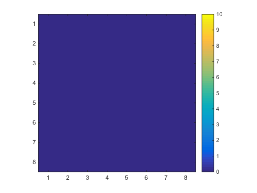 | 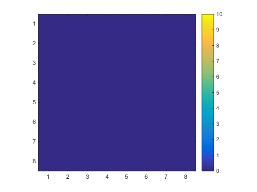 | 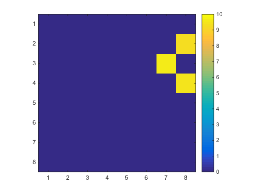 | 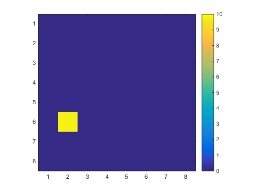 | 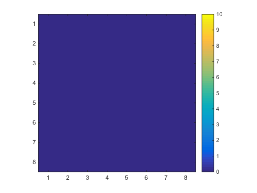 | 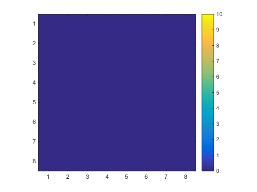 |
|  |  | beta value | 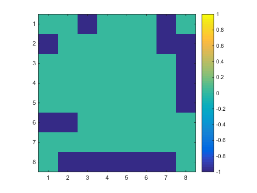 | 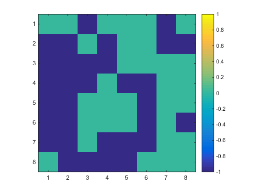 | 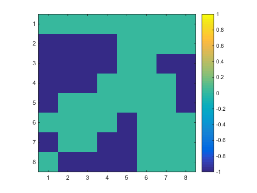 | 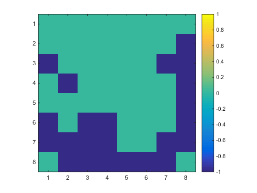 | 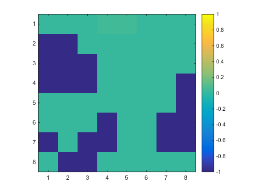 | 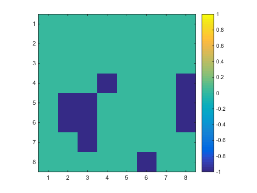 | 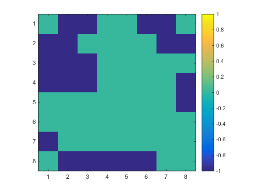 | 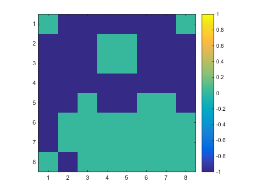 |
|  | Rel | beta choice freq.  (0-10) | 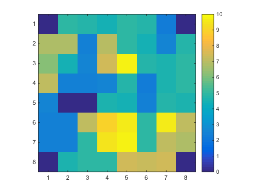 | 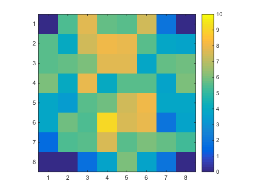 | 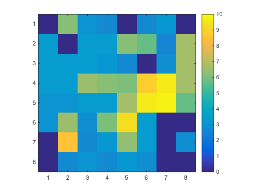 | 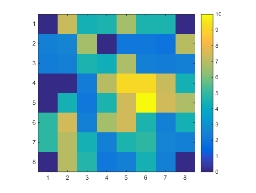 | 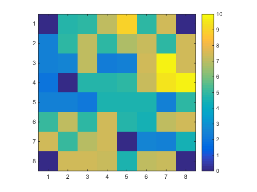 | 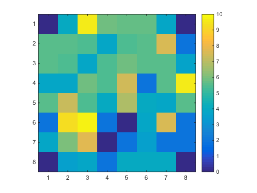 | 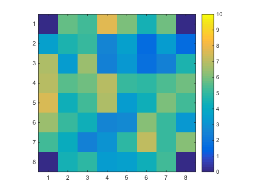 | 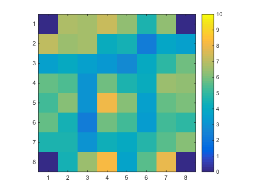 |
|  |  | beta choice freq. >90% | 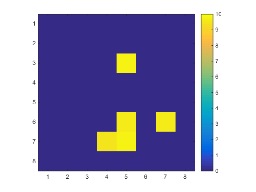 | 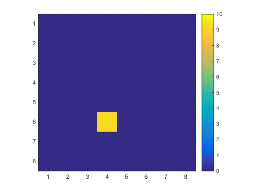 | 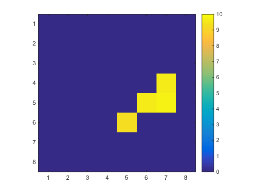 | 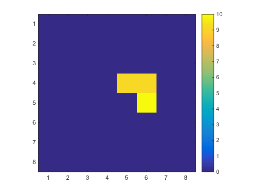 | 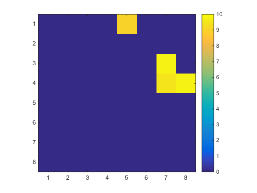 | 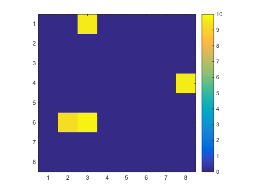 | 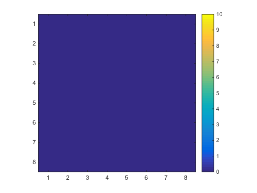 | 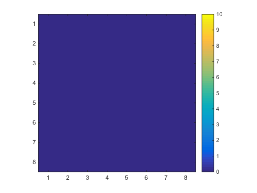 |
|  |  | beta value | 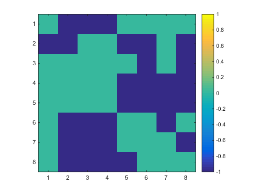 | 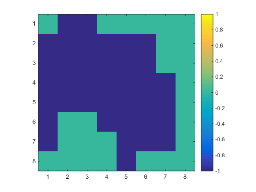 | 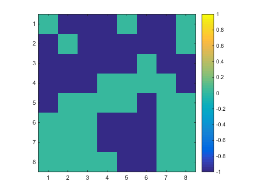 | 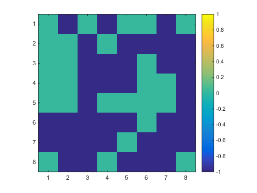 | 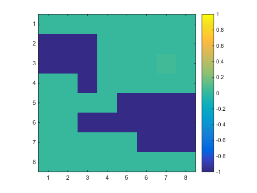 | 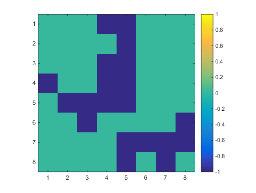 | 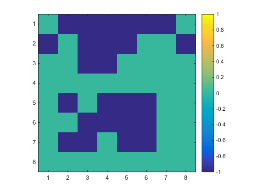 | 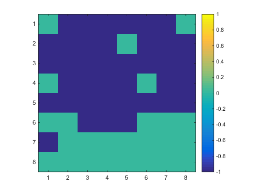 |
| Relatives vs. Controls | Abs | beta choice freq.  (0-10) | 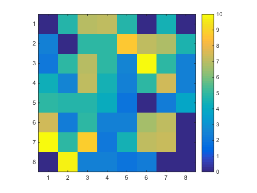 | 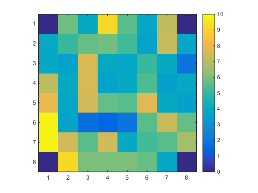 | 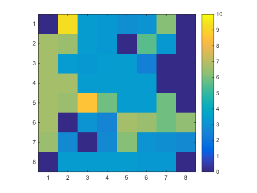 |  |  |  |  |  |
|  |  | beta choice freq. >90% |  |  |  |  |  |  |  |  |
|  |  | beta value |  |  |  |  |  |  |  |  |
|  | Rel | beta choice freq.  (0-10) |  |  |  |  |  |  |  |  |
|  |  | beta choice freq. >90% |  |  |  |  |  |  |  |  |
|  |  | beta value |  |  |  |  |  |  |  |  |
| Age-matched ADHD vs. Relatives | Abs | beta choice freq.  (0-10) |  |  |  |  |  |  |  |  |
|  |  | beta choice freq. >90% |  |  |  |  |  |  |  |  |
|  |  | beta value |  |  |  |  |  |  |  |  |
|  | Rel | beta choice freq.  (0-10) |  |  |  |  |  |  |  |  |
|  |  | beta choice freq. >90% |  |  |  |  |  |  |  |  |
|  |  | beta value |  |  |  |  |  |  |  |  |
| Age-matched Relatives vs. Controls | Abs | beta choice freq.  (0-10) |  |  |  |  |  |  |  |  |
|  |  | beta choice freq. >90% |  |  |  |  |  |  |  |  |
|  |  | beta value |  |  |  |  |  |  |  |  |
|  | Rel | beta choice freq.  (0-10) |  |  |  |  |  |  |  |  |
|  |  | beta choice freq. >90% |  |  |  |  |  |  |  |  |
|  |  | beta value |  |  |  |  |  |  |  |  |

*Note*. Abs = absolute power, Rel = relative power, freq = frequency.

**Table S5.** Beta choice frequencies (all, >90%) and beta values for all the eyes-closed classification models. In beta choice frequency plots the scale 0-10 runs from 0=blue to 10=yellow. In beta choice frequency plots >90% yellow denotes frequency-area features over 90%, blue below 90%. In beta value plots green denotes an increase in power and blue a decrease in power in that scalp area to be a predictor for ADHD status in ADHD vs. Controls models. Similarly, green denotes an increase in power and blue a decrease in power in that scalp area to be a predictor for ADHD status in ADHD vs. Relatives model. Finally, green denotes an increase in power and blue a decrease in power in that scalp area to be a predictor for first-degree relative status in Relatives vs. Controls models. These apply also to the age-matched models.

| **Model** |  |  | **Delta** | **Theta** | **low-Alpha** | **high-Alpha** | **low-Beta** | **mid-Beta** | **high-Beta** | **Gamma** |
| --- | --- | --- | --- | --- | --- | --- | --- | --- | --- | --- |
| ADHD vs. Controls | Abs | beta choice freq.  (0-10) |  |  |  |  |  |  |  |  |
|  |  | beta choice freq. >90% |  |  |  |  |  |  |  |  |
|  |  | beta value |  |  |  |  |  |  |  |  |
|  | Rel | beta choice freq.  (0-10) |  |  |  |  |  |  |  |  |
|  |  | beta choice freq. >90% |  |  |  |  |  |  |  |  |
|  |  | beta value |  |  |  |  |  |  |  |  |
| ADHD vs. Relatives | Rel | beta choice freq.  (0-10) |  |  |  |  |  |  |  |  |
|  |  | beta choice freq. >90% |  |  |  |  |  |  |  |  |
|  |  | beta value |  |  |  |  |  |  |  |  |
| Relatives vs. Controls | Abs | beta choice freq.  (0-10) |  |  |  |  |  |  |  |  |
|  |  | beta choice freq. >90% |  |  |  |  |  |  |  |  |
|  |  | beta value |  |  |  |  |  |  |  |  |
|  | Rel | beta choice freq.  (0-10) |  |  |  |  |  |  |  |  |
|  |  | beta choice freq. >90% |  |  |  |  |  |  |  |  |
|  |  | beta value |  |  |  |  |  |  |  |  |
| matched ADHD vs. Relatives | Rel | beta choice freq.  (0-10) |  |  |  |  |  |  |  |  |
|  |  | beta choice freq. >90% |  |  |  |  |  |  |  |  |
|  |  | beta value |  |  |  |  |  |  |  |  |

*Note*. Abs = absolute power, Rel = relative power, freq = frequency.

**Table S6.** Beta choice frequencies (all, >90%) and beta values for the eyes-closed and eyes-open theta/beta ratio (TBR) classification models. In beta choice frequency plots the scale 0-10 runs from 0=blue to 10=yellow. In beta value plots the scale -.1-.1 runs from -.1=blue to .1=yellow. In beta value plots higher (=yellow) values denote an increase in TBR and lower (=blue) values denote a decrease in TBR in that scalp area to be a predictor for ADHD status in ADHD vs. Controls models. Similarly, higher (=yellow) values denote an increase in TBR and lower (=blue) values denote a decrease in TBR in that scalp area to be a predictor for ADHD status in ADHD vs. Relatives model. Finally, higher (=yellow) values denote an increase in TBR and lower (=blue) values denote a decrease in TBR in that scalp area to be a predictor for first-degree relative status in Relatives vs. Controls models. These apply also to the age-matched models.

| **Model** | **RS condition** | **Sig.** |  | **all** | **age-matched** |
| --- | --- | --- | --- | --- | --- |
| ADHD vs. Controls | TBR eyes-open | ns. | beta choice freq.  (0-10) |  | NA |
|  |  |  | beta value  (-.1-.1) |  | NA |
|  | TBR eyes-closed | ns. | beta choice freq.  (0-10) |  | NA |
|  |  |  | beta value  (-.1-.1) |  | NA |
| ADHD vs. Relatives | TBR eyes-open | ns. | beta choice freq.  (0-10) |  |  |
|  |  |  | beta value  (-.1-.1) |  |  |
|  | TBR eyes-closed | all ns. but age-matched sig | beta choice freq.  (0-10) |  |  |
|  |  |  | beta value  (-.1-.1) |  |  |
| Relatives vs. Controls | TBR eyes-open | all sig. but age-matched ns. | beta choice freq.  (0-10) |  |  |
|  |  |  | beta value  (-.1-.1) |  |  |
|  | TBR eyes-closed | sig. | beta choice freq.  (0-10) |  |  |
|  |  |  | beta value  (-.1-.1) |  |  |

*Note*. TBR = theta/beta ratio, sig. = statistically significant, freq. = frequency.

**Video legends**

**Video 1.** Beta choice frequency of spatial features across all 45 frequencies for eyes-open absolute power in ADHD vs. control classification model.

**Video 2.** Beta choice frequency of spatial features across all 45 frequencies for eyes-open relative power in ADHD vs. control classification model.

**Video 3.** Beta choice frequency of spatial features across all 45 frequencies for eyes-closed absolute power in ADHD vs. control classification model.

**Video 4.** Beta choice frequency of spatial features across all 45 frequencies for eyes-closed relative power in ADHD vs. control classification model.

**Video 5.** Beta choice frequency of spatial features across all 45 frequencies for eyes-open absolute power in first-degree relative vs. control classification model.

**Video 6.** Beta choice frequency of spatial features across all 45 frequencies for eyes-open relative power in first-degree relative vs. control classification model.

**Video 7.** Beta choice frequency of spatial features across all 45 frequencies for eyes-closed absolute power in first-degree relative vs. control classification model.

**Video 8.** Beta choice frequency of spatial features across all 45 frequencies for eyes-closed relative power in first-degree relative vs. control classification model.

**Video 9.** Beta choice frequency of spatial features across all 45 frequencies for eyes-open absolute power in ADHD vs. first-degree relative classification model.

**Video 10.** Beta choice frequency of spatial features across all 45 frequencies for eyes-open relative power in ADHD vs. first-degree relative classification model.

**Video 11.** Beta choice frequency of spatial features across all 45 frequencies for eyes-open absolute power in first-degree relative vs. control classification model with age-matched groups.

**Video 12.** Beta choice frequency of spatial features across all 45 frequencies for eyes-open relative power in first-degree relative vs. control classification model with age-matched groups.

**Video 13.** Beta choice frequency of spatial features across all 45 frequencies for eyes-open absolute power in ADHD vs. first-degree relative classification model with age-matched groups.

**Video 14.** Beta choice frequency of spatial features across all 45 frequencies for eyes-open relative power in ADHD vs. first-degree relative classification model with age-matched groups.

**Video 15.** Beta choice frequency of spatial features across all 45 frequencies for eyes-closed relative power in ADHD vs. first-degree relative classification model with age-matched groups.

**Supplementary Method 1.** Exclusion and inclusion criteria for the ADHD, first-degree relative and control participants.

Exclusion criteria for all participants included: being under 18, history of traumatic brain injury (e.g. concussion), medical conditions such as epilepsy, severe migraine, hearing and/or severe motor impairment, stroke and diabetes, psychiatric conditions such as psychosis, bipolar disorder, eating disorder, schizophrenia and personality disorder, learning and/or intellectual disability, history of alcohol or drug use disorder, current medication use of antidepressants (e.g. selective serotonin reuptake inhibitors (SSRIs)), benzodiazepines, antipsychotics or anticonvulsants. Furthermore, ADHD group inclusion criteria required the participant to have a self-reported formal diagnosis of ADHD, a T score over 65 in the ADHD index subscale of Conners’ Adult ADHD Rating Scale (CAARS) and willingness to abstain from medication 36 hours prior to testing (letter detailing abstinence was sent to their GP). The first-degree relative group inclusion criteria required the relative to share both biological parents or to be the biological parent of a person diagnosed with ADHD. For inclusion in the healthy control group, absence of ADHD was required (i.e. no diagnosis of ADHD and a T score below 65 in the ADHD index subscale of CAARS).

On the day of data collection, participants were screened for co-morbidities using a shortened version of the Structural Clinical Interview DSM-IV (SCID; including subsections screening for psychosis, mood disorders, dysthymia, bipolar disorder, suicidality, eating disorders and substance abuse).

**Supplementary Method 2.** Logistic regression machine learning analysis with the Regularized Adaptive Feature Thresholding algorithm (RAFT) and Elastic Net

*1. Nested cross-validation*

The dataset is initially divided into 10 cross-validation (CV) folds. The entire analysis is performed 10 times, using 90% of the dataset (the training set) to create a regression model which is then tested on the remaining 10% of the data (the test set). In each subfold (inner cross-validation), the data were z-scored and extreme values were replaced with a value of 3 (i.e., Winsorizing). Within the training set, additional ‘nested’ cross-validation with 10 partitions is used to support the analyses at the feature selection and model optimisation level. Results from all 10 CV folds are finally aggregated, using the frequency with which a variable is found in models from different CV folds as a measure of its robustness.

*2. Feature thresholding*

Each feature of the dataset is individually evaluated to assess its utility in predicting the discrete target variable (i.e. group membership), in what can be considered a filtering step. A simple logistic regression model is applied to the training set (81% of the data) for that feature and the target variable, and the resulting regression weight is used to make outcome predictions for the test set (9% of the data). The model was evaluated on the nested test set and the model’s accuracy was measured with the Area Under the Curve (*AUC*). AUC was calculated using fastAUC function (www.mathworks.com/matlabcentral/fileexchange/50962-fast-auc) in MATLAB.

Based on the range of AUC values estimated across all CV folds, a set of ten *prediction error* *thresholds* is created. These thresholds rank features according to their AUC. At each prediction error threshold (*t_AUC_*) and for each nested CV partition *n* in every main CV fold *m* there is a subset of features *f* which have smaller prediction error values than that threshold.

$$s_{m,n}\left( t_{AUC} \right)=\left\{ f \right| {AUC}_{m,n}\left( f \right)<t_{mse}\}$$

A set of ten *stability* *thresholds* (t_stability_) is also used to assess how stable, over samples, the AUC value assigned to each feature is. This is quantified as the number of nested CV partitions *n* in which a feature had a prediction error value lower than each AUC threshold.

$${stability}_{m, t_{AUC}}\left( f \right)= \left| \left\{ n \right|f\in s_{m,n}\left( t_{AUC} \right)\} \right|$$

The ten prediction error thresholds and ten stability thresholds jointly define 100 new summary datasets, which include all features that had a smaller AUC value than t_AUC_ in the number of CV partitions specified by t_stability_.

$$D_{m}\left( t_{AUC}, t_{stability} \right)=\left\{ f \right| {stability}_{m,t_{AUC}}(f)\geq t_{stability}\}$$

The prediction error thresholds are chosen based on the range of prediction error and feature stability across the sample. The most liberal prediction error threshold *t_AUC_(max)* is chosen such that in each main CV fold *m* there is at least one feature which is common to all nested CV partitions, i.e. t_AUC_(max) is the smallest prediction error value at which the following is true:

$${|\{m |D_{m}(t}_{AUC}\left( \max\right),10)\neq\emptyset\}|=10$$

The strictest prediction error threshold *t_AUC_(min)* is set as the lowest possible prediction error value at which every nested CV partition *n* in every main CV fold still contains at least one feature that has a smaller prediction error value than that threshold. That is, *t_AUC_(min)* is the smallest prediction error value at which the following is true:

$$|{\left\{ (m,n) \right| s}_{m,n}(t_{AUC}\left( min \right))\neq\emptyset\}|=100$$

Taken together *t_AUC_* and *t_stability_* define how high the individual predictive power of each feature in the knowledge base is, and how stable results with each feature are across subsets of the sample. The creation of these thresholds serves the purpose of integrating the choice of the criterion used to select features from the filtering step into model selection, eliminating researcher input at this point.

*3. Model Optimisation*

The feature sets which are created in the first analysis step are used as inputs into a model optimisation algorithm. We chose to use Elastic Net regularisation (1-2), which is a regularization method for generalized linear models that includes regularization (i.e., lasso regularization 𝑙1 - least absolute shrinkage and selection operator) and 𝑙2 regularization (as in ridge regularization). Lasso regularization allows parameters to be 0, promoting parsimonious solutions; whereas ridge regularization allows parameters to be small but not to reach 0, avoiding overfitting (3). The Elastic Net uses complexity and weighting parameters (𝜆 and 𝛼, respectively) that are not known a priori. Therefore, a range of values were explored: 20 linearly-spaced values of both parameters in the range of 0.01 to 1 and all their possible combinations. Both Lasso and Ridge regression apply a penalty for large regression coefficient values, but Lasso regularization favours models with fewer features, making it more prone to excluding features. The prediction accuracy of each parameter combination was assessed using the mean squared error. The parameter combination that yielded the lowest error was selected per subfold. The mode of 𝛼 and the median of 𝜆 across subfolds were selected as parameters per main fold.

$$d_{m,n}\left( t_{AUC}, t_{stability}, \alpha, \lambda\right)\subseteq D_{m}\left( t_{AUC}, t_{stability} \right)$$

*4. Model validation*

After the model optimisation step, the combination of model parameters and thresholds which resulted in the model with the lowest prediction error is identified for each nested CV partition. The optimal model parameters and thresholds from each nested CV partition are used to identify what parameters will be used to create the final prediction model in each main CV fold, using the most frequently occurring values of *α*, *λ*, *t_AUC_* and *t_stability_*. To select the features to include in the final model for each CV fold, the stability of all features that were included in the updated feature sets at the optimal prediction error and stability thresholds is re-calculated.

$${stability}_{m}\left( f \right)= \left| \left\{ n \right|f\in d_{m,n}\left( t_{AUC}, t_{stability}, \alpha, \lambda\right)\} \right|$$

Only the features that were included in at least as many of the ten models with optimal parameters as specified by the optimal stability threshold are used to create the feature set for the final model.

$${FeatureSet}_{m}=\left\{ f \right| {stability}_{m}(f)\geq t_{stability}\}$$

It is possible that this implementation of the stability threshold does not leave any features for inclusion in the model. Should this be the case the closest possible parameter combination is used to create the feature set.

This feature set and the covariates are used as input into the Elastic Net, using the optimal values for *α* and *λ*, and the entire training set (90% of the data). The beta weights generated by the Elastic Net are subsequently used to make outcome prediction for the final unseen portion of the data (10%). Each CV fold is used to make outcome predictions for 10% of the data, and the evaluation of model fit is carried out using the complete vector of outcome predictions from all CV folds. The prediction of the model on the test set of each main fold was saved and pooled across main folds.

Performing model selection as an integrated step within the CV framework is an essential step in preserving the external validity of the resultant model (4).

**Schematic description of the model optimization part in the logistic regression machine learning analysis procedure used.**

**References (Supplementary Method 2)**

1. Friedman J, Hastie T, Tibshirani R. Regularization paths for generalized linear models via coordinate descent. 2010. *J Stat Softw*. 33(1):1.

2. Zou H, Hastie T. Regularization and variable selection via the elastic net. 2005. *J R Stat Soc Series B Stat Methodol*. 67(2):301-20.

3. Murphy, D., Spooren, W. EU-AIMS: A boost to autism research*. Nat Rev Drug Discov*. 2012;11:815–816.

4. Cawley GC, Talbot NL. On over-fitting in model selection and subsequent selection bias in performance evaluation. 2010. *J Mach Learn Res*. 11:2079-107.
